# Estimating trial-wise modulation of functional connectivity using event-related fMRI

**DOI:** 10.64898/2026.07.15.738746

**Authors:** Kai Hwang, Shannon E. Stokes, Stephanie C. Leach, Jiefeng Jiang

## Abstract

Understanding the neural basis of human cognition requires measuring not only localized brain activity but also how functional interactions between brain regions change in response to different cognitive demands. Event-related fMRI is an efficient design for linking trial-wise behavioral and computational variables to brain activity, but comparable methods for examining their effects on functional connectivity remain limited. Here, we introduce a regression-based beta-series modeling (BSM) framework that leverages single-trial response estimates from event-related fMRI to quantify how task-related variables modulate functional connectivity. By modeling the interaction between regional activity and task-related variables, this framework provides a statistical approach for testing connectivity modulation by both continuous and categorical variables. We evaluated this approach using comprehensive simulations across several experimental conditions and signal qualities. The regression-based BSM approach can sensitively detect ground-truth effects with well-controlled false-positive rates. Compared with generalized psychophysiological interaction, BSM achieved comparable performance across most conditions while demonstrating improved statistical power under lower signal-to-noise conditions. We further validated the approach using empirical event-related fMRI data. Distinct trial-wise cognitive variables selectively modulated functional connectivity during their corresponding trial epochs, demonstrating the temporal specificity and flexibility of the approach. Together, these findings demonstrate that regression-based BSM provides a flexible framework for examining task-modulated functional connectivity in event-related fMRI, with applications ranging from conventional experimental manipulations to continuous computational model-derived variables.

## 1. Introduction

Human cognition emerges from interactions among brain regions rather than isolated activity (Goldman-Rakic, 1988; Mesulam, 1990). Distributed yet coordinated neural activity supports diverse cognitive functions, including perception, decision making, learning, and cognitive control (Cole et al., 2013; Hipp et al., 2011; van den Brink et al., 2022). Understanding how brain network connectivity is configured in response to different cognitive demands is therefore a major goal of cognitive neuroscience. Functional connectivity analyses provide a useful framework for addressing this question. By quantifying statistical dependencies between neural signals (Park & Friston, 2013), functional connectivity assesses the extent activity in different brain regions covaries over time. These statistical relationships provide a network level description of how distributed brain systems are organized. Unlike localization analyses, which identify where task-evoked responses occur, functional connectivity characterizes how activity is coordinated across distributed brain regions. Importantly, these interactions have been shown to change across behavioral contexts (Cocuzza et al., 2020; Hearne et al., 2017). As task demands change, the strength of coupling between regions may increase or decrease, reflecting dynamic reconfiguration of region-wise interactions. This approach naturally emphasizes that brain networks are not static, but instead dynamically reconfigure in response to different task demands.

Although the task-dependent and dynamic nature of brain network interactions have long been recognized (Mill et al., 2017; Shine et al., 2016), much of the work has focused on task-free, resting-state connectivity (Buckner et al., 2013; Deco et al., 2011). These studies have provided insights into the organization of large-scale functional brain networks (Ladwig et al., 2026; Schaefer et al., 2018), and its alterations across neurological and psychiatric disorders (Greicius et al., 2004; Yan et al., 2019). However, resting-state connectivity does not directly address how functional connectivity are reconfigured in response to changing cognitive demands. For this purpose, methods that quantify connectivity during task performance are needed.

Several methods have been developed to study task-dependent changes in functional connectivity. For example, Psychophysiological interaction (PPI) analysis tests whether connectivity between brain regions varies as a function of an experimental manipulation (Friston et al., 1997). Generalized PPI extends this framework to accommodate multiple task conditions in event-related fMRI (McLaren et al., 2012). A different approach is beta-series correlation (Rissman et al., 2004), which estimates trial-wise response amplitudes and measures connectivity by correlating beta estimates across brain regions. In conventional beta-series approaches, task-modulated connectivity is often assessed by estimating connectivity separately for different experimental conditions and comparing the resulting correlation coefficients. Beyond comparing condition-specific correlations, task-modulated connectivity can also be estimated using a regression model that includes regional beta-series activity, a task variable, and their interaction. Di et al. (2021) examined this approach using a categorical task contrast and demonstrated its relationship to conventional PPI and beta-series correlation approaches. In this framework, the interaction coefficient directly tests whether the relationship between source and target activity varies with the task variable. The same regression approach can also accommodate continuous moderators, allowing connectivity modulation to be estimated as a function of trial-wise behavioral or computational variables rather than discrete experimental conditions. Both PPI and beta-series approaches explicitly model task-evoked responses to quantify task-dependent changes in connectivity. In contrast, residual-based functional connectivity approaches estimate task connectivity after removing trial-evoked responses from continuous BOLD time series (Al-Aidroos et al., 2012; Hwang et al., 2019). Extensive work has been conducted to compare and evaluate different task connectivity methods (Cisler et al., 2014; Cole et al., 2019; Di et al., 2021; Masharipov et al., 2024; O’Reilly et al., 2012).

Task-based connectivity methods aim to characterize how interactions between brain regions support different cognitive processes. Cognitive processes are inherently dynamic and flexible, and event-related fMRI provides an efficient experimental design approach for isolating these dynamics (Buckner, 1998). It enables trial-by-trial manipulations of cognitive processes, as well as temporally separating distinct cognitive operations between epochs within a trial (Ollinger et al., 2001), such as stimulus input, decision making, response selection, and feedback processing. Event-related fMRI design thus allows trial-level variation in cognition to be quantified at the trial and epoch level. A major challenge is therefore how to characterize this trial- and epoch-specific cognitive process and its variability. One useful approach is computational modeling, which estimates latent variables that evolve throughout a task to provide mechanistic inferences of underlying cognitive processes. Examples include prediction errors (Gläscher et al., 2010), expected value (Daw et al., 2011), uncertainty (Nassar et al., 2019), latent-state estimates (Schuck et al., 2016), and memory strength (Li et al., 2025). These variables are typically continuous and change trial to trial, and can be used to model regional BOLD activity to test how computational variables are instantiated in the brain. An equally important question is whether these variables also influence interactions between distributed brain regions.

Addressing this question requires a statistical framework that can directly estimate how functional connectivity varies with both continuous and categorical task-related variables.

Existing task-connectivity approaches differ in how they accommodate trial-wise variation and estimate task-dependent changes in connectivity. Residual-based methods require continuous time series after removal of task-evoked responses. Consequently, they are better suited for studying sustained states of connectivity but not trial-to-trial variation. PPI approaches are better suited for this goal because they can model interactions between a physiological signal and a psychological variable. However, PPI was originally developed to test categorical task conditions as psychological regressors. Although continuous parametric modulators can be incorporated into PPI models, it remains unclear whether this approach provides sensitive estimates for rapidly varying trial-level and epoch-specific variables commonly generated in event-related experiments, including computational model-derived variables. Beta-series methods provide a complementary approach because they estimate regional responses at the level of individual trials. By applying a moderation regression to these trial-wise estimates, task-dependent connectivity can be quantified directly through an interaction coefficient, rather than through separate condition-specific correlations. This approach extends the flexibility of beta-series connectivity analyses while retaining their compatibility with event-related experimental designs.

Here, we systematically evaluate a regression-based beta-series modeling (BSM) framework for estimating task-modulated functional connectivity in event-related fMRI. Building on previous work applying interaction regression to beta-series connectivity (Di et al., 2021), this approach models how trial-wise variables moderate the relationship between regional response amplitudes. Single-trial response amplitudes are first estimated using least-squares separate (LSS) regression (Mumford et al., 2012). Connectivity is then modeled across trial-wise response amplitudes rather than continuous BOLD time series. The interaction coefficient quantifies whether functional connectivity systematically changes with the moderating variable. We evaluate the statistical properties of this framework and demonstrate its application to continuous, computationally derived trial-wise variables.

An initial application of this approach was included in a previous empirical study (Leach et al., 2026), where we used it to investigate task-related changes in frontoparietal functional connectivity. The present study extends this initial application by systematically evaluating the statistical properties and performance of regression-based BSM. We first characterized its statistical properties using simulations to examine parameter recovery, statistical power, false-positive control, signal-to-noise ratio (SNR), and robustness to intrinsic coupling and task-evoked response coactivation. We also compared this method with PPI across a broad range of SNR and experimental conditions. We then validated the method using an event-related fMRI dataset in which trial-wise computational variables were obtained from a computational model fitted to human behavioral data (Leach et al., 2025, 2026). Together, these analyses showed BSM as a practical method for investigating functional connectivity in event-related fMRI experiments.

## 2. Methods

### 2.1 Overview

The BSM method quantifies whether functional connectivity between two brain regions is modulated by a trial-wise variable. Similar to conventional seed-based functional connectivity analyses, one brain region is designated as the seed region, and functional connectivity is modeled as the statistical relationship between trial-wise response amplitudes in the seed and target brain regions. The method then tests whether this relationship changes systematically as a function of a trial-wise moderating variable.

BSM separates three sources that contribute to task-related functional connectivity estimates. First, baseline coupling represents the relationship between source and target activity that is independent of the computational variable and task-evoked responses. This is akin to task-free, intrinsic functional connectivity (Cole et al., 2014; Fox et al., 2006). Second, task-evoked responses capture changes in local activity magnitudes that are evoked by task inputs (e.g., computational variable or stimulus conditions) and thus do not reflect changes in functional connectivity (O’Reilly et al., 2012). Third, connectivity modulation represents the extent to which the computational variable modulates functional connectivity between the source and target regions. This serves as the primary measurement of interest. The following sections describe the mathematical formulation of the BSM model, the simulation framework we used to evaluate its statistical properties, and how we validated it with an empirical event-related fMRI dataset.

### 2.2 Mathematical formulation of BSM

Let *S_i_* denote a source region’ activity on trial *i*, *T_i_* denote target region’s activity, and *M_i_* denote a trial-wise moderator variable derived from a computational model. Connectivity modulation is estimated using a regression model:

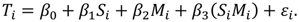

The intercept term *β_0_* captures the mean response in the target region. The coefficient *β_1_* represents baseline coupling between source and target activity. This term is conceptually identical to intrinsic connectivity independent of task-evoked responses. The coefficient β_2_ captures how strong the moderator elicits task-related activity in the target region, akin to task-evoked responses. We will refer to β_2_ as moderator main effect for the rest of the manuscript. The interaction coefficient β_3_ quantifies the degree to which the moderator influences connectivity between source and target activity and is the connectivity modulation effect of interest (Fig 1).

**Fig 1.**
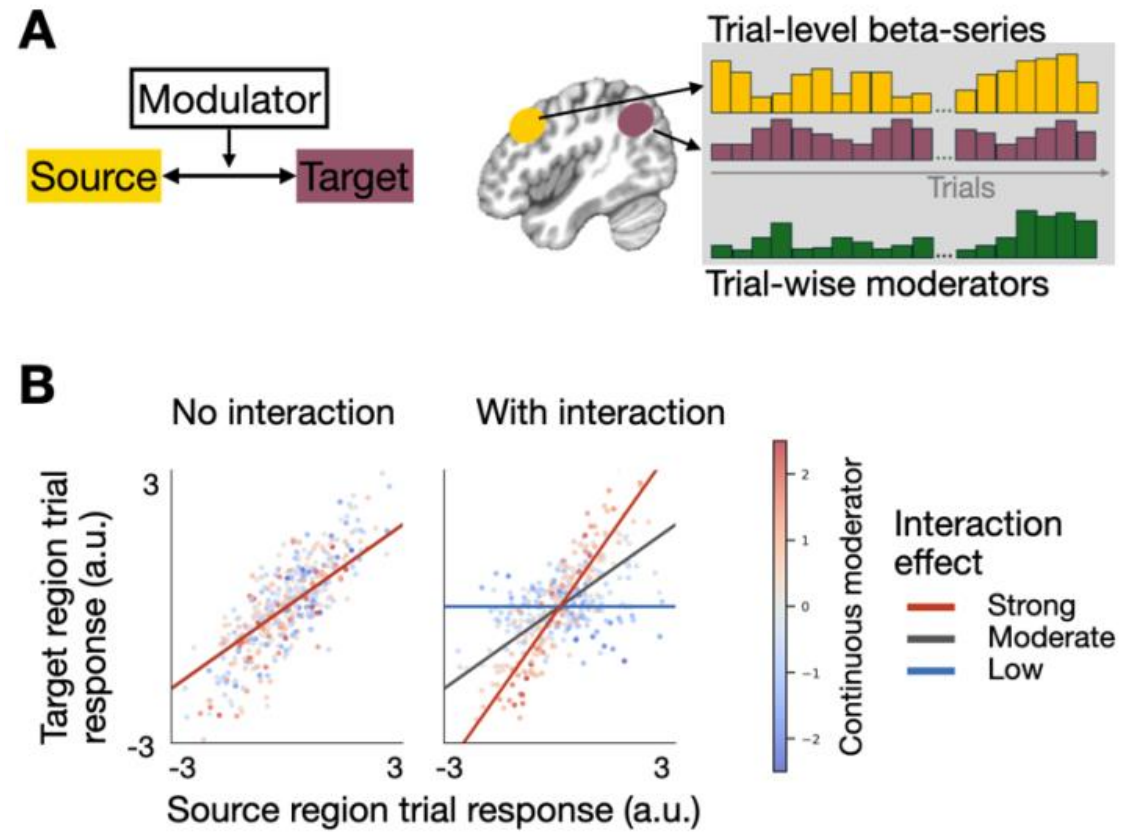
Conceptual illustration of BSM. (A) The overall principle of BSM, where activity estimates were extracted from a source and target brain regions to form a beta-series, and its interaction with a trial-wise moderator can be tested in a regression model. (B) Trial-wise source and target responses were simulated to illustrate the statistical principle underlying BSM. Each point represents the response amplitude from a single trial, analogous to single-trial β estimates obtained from LSS analyses. Point color indicates the value of a continuous trial-wise moderator, with cooler and warmer colors corresponding to lower and higher moderator values, respectively. Colored lines illustrate how the moderator can exert different levels of interaction effect on the coupling between source and target activity.

This implementation can be interpreted as a trial-wise version of the classical psychophysiological interaction analysis (Friston et al., 1997). However, because estimation occurs at the level of single-trial activity amplitudes, the method is suited for trial-by-trial variables and rapid event-related design.

### 2.3 Simulations

We performed a series of simulations to characterize the statistical properties of regression-based BSM under realistic event-related fMRI conditions. Although the statistical properties of interaction under regression theory are well established, applying the BSM framework to fMRI requires estimating single-trial response amplitudes from noisy BOLD signals. Variability in hemodynamic responses, temporal overlap between events, and noise can introduce errors in single-trial estimates that propagate into subsequent connectivity analyses. Our simulations therefore aimed to evaluate the performance of the complete BSM estimation pipeline when applied to event-related fMRI data, focusing on parameter recovery, statistical power, and false-positive rates.

We simulated trial-wise latent neural activity and functional connectivity between a source region and a target region using a generative model with known parameters. These latent signals were transformed into BOLD responses using a hemodynamic forward model that incorporated variability in hemodynamic responses and temporally autocorrelated noise. Single-trial response amplitudes were then estimated using least-squares separate (LSS) regression and entered into the BSM model to estimate connectivity modulation. This approach allowed us to evaluate how well BSM recovered known interaction effects from simulated BOLD data rather than directly from the latent neural signals.

We systematically varied interaction strength, baseline coupling, task-evoked source activity, latent neural signal-to-noise ratio (SNR), and BOLD SNR to characterize their influence on interaction estimates and statistical inference. Additional simulations independently manipulated the correlation between source activity and the moderator to examine its effects on interaction estimation, statistical power, and false-positive rates. We also varied experimental design parameters, including the number of trials, sample size, repetition time, and inter-trial interval, to assess their influence on statistical sensitivity. Finally, we compared the performance of BSM with a conventional PPI approach using the same simulated BOLD data.

#### 2.3.1 Event-related design and latent generative model

Simulations were designed to follow common event-related fMRI designs. For each simulation, trial onset times were generated by sampling inter-trial intervals (ITIs) from an exponential distribution with a specified mean. To avoid overly short intervals, ITIs were constrained to a minimum duration of 0.5 second. Each trial was modeled as a brief event with a simulated duration. BOLD signals were sampled at a repetition time (TR) of 2 second. Later, we also explored the effects of different TR lengths and stimulus durations.

For each simulated subject, trial-wise latent neural activity was generated for a source region and a target region. Latent signals simulations were designed to separate three sources that can influence task-based connectivity: (1) co-activation of task-evoked responses, (2) intrinsic, baseline functional coupling without task inputs, and (3) modulation of functional connectivity by a trial-wise variable.

On each trial *i*, a moderator variable *M_i_* was sampled from a standard normal distribution and standardized to have zero mean and unit variance. The moderator represents a generic model-derived variable, such as uncertainty, belief strength, or prediction error. Latent activity in the source region was generated according to

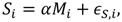

where *S_i_* denotes source region activity, *α* controls the magnitude of the evoked activity of *M_i_* effect, and *ε_S_*_,*i*_, represents the noise. This formulation produces a source region whose activity covaries with the moderator. To simulate task-modulated connectivity, an interaction drive was defined as

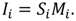

and the target region’s activity was then generated according to

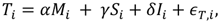

where *T_i_* denotes target region activity, *γ* controls baseline coupling between source and target regions, *δ* controls the magnitude of connectivity modulation, and *ε_T_*_,*i*_ represents independent noise. The term *γS_i_* produces intrinsic coupling that is independent of the moderator, whereas the term *αM_i_* introduces co-activated task-evoked responses in both regions. For simplicity, we assume task-evoked response is the same for both source and target regions. The interaction term *δI_i_* represents the ground truth connectivity modulation effect and constitutes the primary parameter of interest throughout the simulations.

After generating latent signals, we further manipulated the latent neural signal-to-noise ratio (SNR). This manipulation was included because the fidelity with which a computational variable is represented in neural activity is expected to vary across brain regions, experiments, and participants (e.g., subject variability). For both source and target regions, latent neural noise was sampled independently from a normal distribution and standardized to have zero mean and unit variance. Noise magnitude was then scaled relative to the standard deviation of the signal component to achieve a desired latent SNR. Specifically, latent activity was generated according to

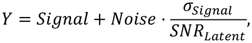

where *σ_Signal_* denotes the standard deviation of the signal component and *SNR_Latent_* specifies the desired latent neural SNR.

To examine whether correlations among regression predictors influence BSM estimation, we conducted additional simulations in which the correlation between source activity and the moderator was manipulated independently of baseline source–target coupling. Specifically, source activity was generated as

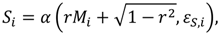

where *r* specifies the source–moderator correlation, and *M*i and *ε_S_*_,*i*_ are independently generated and standardized. We varied *r* across 0, 0.25, 0.50, 0.75, and 0.90. All other simulation parameters were held constant (latent SNR = 0.5, BOLD SNR = 1.0). This simulation allowed us to evaluate the effects of source–moderator correlation and baseline coupling on interaction estimates, statistical power, and false-positive rates.

To assess whether temporal dependencies in the moderator affect BSM estimation, we conducted additional simulations comparing three moderator properties: (1) independent Gaussian values (as described above), (2) autocorrelation generated using a first-order autoregressive process with an autoregressive coefficient of 0.8, and (3) block-like values representing persistent trial-wise variables. In the block-like condition, the moderator alternated between two values, with each value persisting for a randomly selected duration of 10–20 trials. These simulations allowed us to assess whether temporal dependencies in the moderator, combined with overlapping hemodynamic responses and temporally autocorrelated BOLD noise, influence false-positive rates and sensitivity to connectivity modulation.

#### 2.3.2 Hemodynamic forward model

Latent neural activity was then transformed into simulated BOLD signals using a canonical hemodynamic response function (HRF). The HRF was implemented with a double-gamma function consisting of a positive peak response and a delayed undershoot component. Trial-wise neural amplitudes were convolved with the HRF at their corresponding event onset times to generate continuous BOLD time series for both source and target regions. Given known variability in HRF between subjects as well as between brain regions (Handwerker et al., 2004), variability in hemodynamic timing was incorporated into the simulations. For each simulated participant, different HRFs were generated for the source and target regions by varying the latency of the positive peak. Specifically, peak delay parameters were sampled from a 2 second range surrounding the canonical 6 second peak latency. Note that therefore source and target regions will have different HRF properties, in addition to individual differences, and thus introducing variability in the true HRF should introduces realistic model mismatch. Following convolution, simulated BOLD signals were sampled at with the simulated TR rate.

To simulate realistic BOLD signals, temporally autocorrelated noise was then added to source and target BOLD signals. Noise was generated using a first-order autoregressive AR(1) process, *n_t_* = *ρn_t_*_−1_ + *η_t_*, where *ρ* is the autoregressive coefficient. The autoregressive coefficient was fixed at 0.17, based on the averaged empirical observations from our previously published fMRI datasets (Chen et al., 2024; Leach et al., 2025). Noise signals were standardized to have zero mean and unit variance before scaling to achieve a desired BOLD signal-to-noise ratio (SNR). Observed BOLD activity was then normalized according to

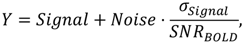

where *σ_Signal_* denotes the standard deviation of the convolved signal component and SNR_BOLD_ specifies the desired SNR.

#### 2.3.3 Single trial response estimate

Single trial response amplitudes were estimated using the LSS regression procedure (Mumford et al., 2012). For each trial j, a design matrix was constructed containing three regressors: (1) a regressor modeling the trial of interest, (2) a regressor modeling all remaining trials combined, and (3) an intercept term. The coefficient associated with the trial-of-interest regressor was retained as the estimated response amplitude for that trial. This procedure will yield one response estimate per trial for both the source and target regions. The resulting beta series therefore provides an estimation of trial-by-trial fluctuations in task-evoked activity while reducing contamination from overlapping responses. However, the accuracy of these estimates can depend on design choices, including the temporal spacing of events and overlap between hemodynamic responses (Abdulrahman & Henson, 2016; Mumford et al., 2014; Visser et al., 2016). Because the BSM framework operates on trial-wise response amplitudes rather than continuous BOLD signals, these beta estimates constitute the inputs to all subsequent connectivity analyses. To assess the fidelity of single-trial response estimation derived from the above procedure, we conducted additional simulations varying the mean ITI (1.5, 3.0, 4.5, and 6.0 s).

Estimation fidelity was quantified separately for the source and target regions using the Pearson correlation between the ground-truth latent trial amplitudes and their corresponding LSS beta estimates.

#### 2.3.4 BSM Connectivity Estimation

Task-modulated connectivity was estimated using the BSM framework described above. For each simulated participant, source and target region’s beta series obtained from the LSS procedure were standardized to have zero mean and unit variance. Then, for each simulated subject, target region’s beta estimates were modeled as a function of source region activity, moderator values, and their interaction. The interaction coefficient served as the subject level estimate of task modulated connectivity. Because source activity and moderator related activation were included explicitly in the regression model, the interaction coefficient reflects variance uniquely attributable to modulation in functional connectivity rather than shared co-activation.

The subject level interaction coefficients were subsequently entered into group-level analyses to assess parameter recovery, statistical power, and false positive rates.

#### 2.3.4 Group-Level Inference

Group-level inference was performed using a one-sample t-test against zero across participants, analogous to a conventional second-level fMRI analysis. For simulation conditions containing a true, non-zero interaction effect, statistical power was quantified as the proportion of simulation repetitions yielding a significant effect at alpha = 0.05. For simulations in which the ground-truth interaction parameter was set to zero, the same procedure was used to estimate the false-positive rate. Parameter recovery was assessed by examining the relationship between the ground-truth interaction parameter and the mean estimated interaction coefficient across simulation conditions.

#### 2.3.5 Simulation parametrization

Parameters set for the latent generative model were varied to explore a range of conditions, including task-evoked source activation (α = 0.1 to 0.75), baseline source-target coupling (γ = 0 to 0.60), connectivity modulation (δ = 0 to 0.40), latent neural signal-to-noise ratio (SNR = 0.1 to 1.0), and BOLD signal-to-noise ratio (SNR = 0.50 to 2.0). Because the true magnitudes of latent neural coupling and task-modulated connectivity are not directly observable in empirical data, these ranges were chosen to span weak through relatively strong effects while avoiding unrealistic ceiling conditions.

#### 2.3.6 Time-series PPI with a parametric psychological regressor

Although both PPI and BSM test whether functional connectivity is modulated by task variables, they differ in several important aspects. PPI estimates interactions in the continuous fMRI time series by first deconvolving the seed-region BOLD signal to approximate latent neural activity, multiplying this neural estimate by the psychological regressor, and convolving the interaction term with a HRF function to form a PPI regressor (McLaren et al., 2012). In contrast, BSM operates directly on trial-specific response estimates obtained from event-related GLMs, testing whether trial-to-trial variation in a moderator variable explains variability in functional connectivity across trials. Consequently, the primary distinction between the two methods lies in whether interactions are estimated from deconvolved estimated time series or from single-trial response estimates. To benchmark BSM against a conventional time-series PPI approach, we implemented PPI with a single parametric psychological regressor using custom Python code. For each simulated participant, the source-region BOLD signal was deconvolved using a discrete cosine transform basis to reconstruct latent neural activity. The resolution was set to upsample 10 times from the BOLD sampling rate. Basis coefficients were estimated using ridge-regularized linear inversion with a regularization parameter of 0.05. The reconstructed neural signal and the psychological regressor were each standardized to zero mean and unit variance before multiplication to obtain the neural interaction term. This interaction term was subsequently convolved with a canonical double-gamma HRF and down sampled to the original BOLD sampling rate. The target-region BOLD signal was then regressed on the source-region BOLD signal, the HRF-convolved psychological regressor, and the PPI interaction regressor, with all regressors standardized before estimation. Subject-level interaction coefficients were entered into the same group-level inference procedure used for BSM. We performed the above simulation both with and without variability in HRF timing (see Methods 2.3.2).

### 2.4 Empirical Validation

To demonstrate the applicability of BSM to empirical fMRI data, we applied the method to a previously published event-related fMRI dataset examining cognitive integration (Leach et al., 2025, 2026). In this experiment, 38 participants integrated probabilistic sensory evidence with an internally maintained state representation to determine whether to perform a face or scene judgment on each trial. Behavioral performance was modeled using a Bayesian model that generated trial-wise estimates of task belief, reflecting the inferred probability that a particular task should be performed given the available sensory and contextual information. Full details regarding participant recruitment, behavioral procedures, computational modeling, MRI data acquisition, and preprocessing are reported in our published studies (Leach et al., 2025, 2026). Below we briefly describe the experimental paradigm and computational model.

The experimental paradigm was designed to require participants to integrate external sensory evidence with an internally maintained contextual representation to determine which task should be performed on each trial (Fig. 2A). Each trial began with a random-dot color display in which participants judged whether red or yellow dots were more dominant (Fig. 2B). The strength of the sensory evidence varied continuously across trials by manipulating the proportion of the dominant color, thereby parametrically varying perceptual uncertainty. After a delay, participants viewed a composite face-scene image and performed either a face judgment (male vs. female) or a scene judgment (city vs. nature landscape) task. Critically, the mapping between cue color and task depended on a hidden contextual state that covertly reversed unpredictably every 10–20 trials (Fig 2C). Thus, successful performance required participants to integrate the sensory evidence with an internally estimated hidden state to infer which task should be performed. Feedback was provided after each response.

**Fig 2.**
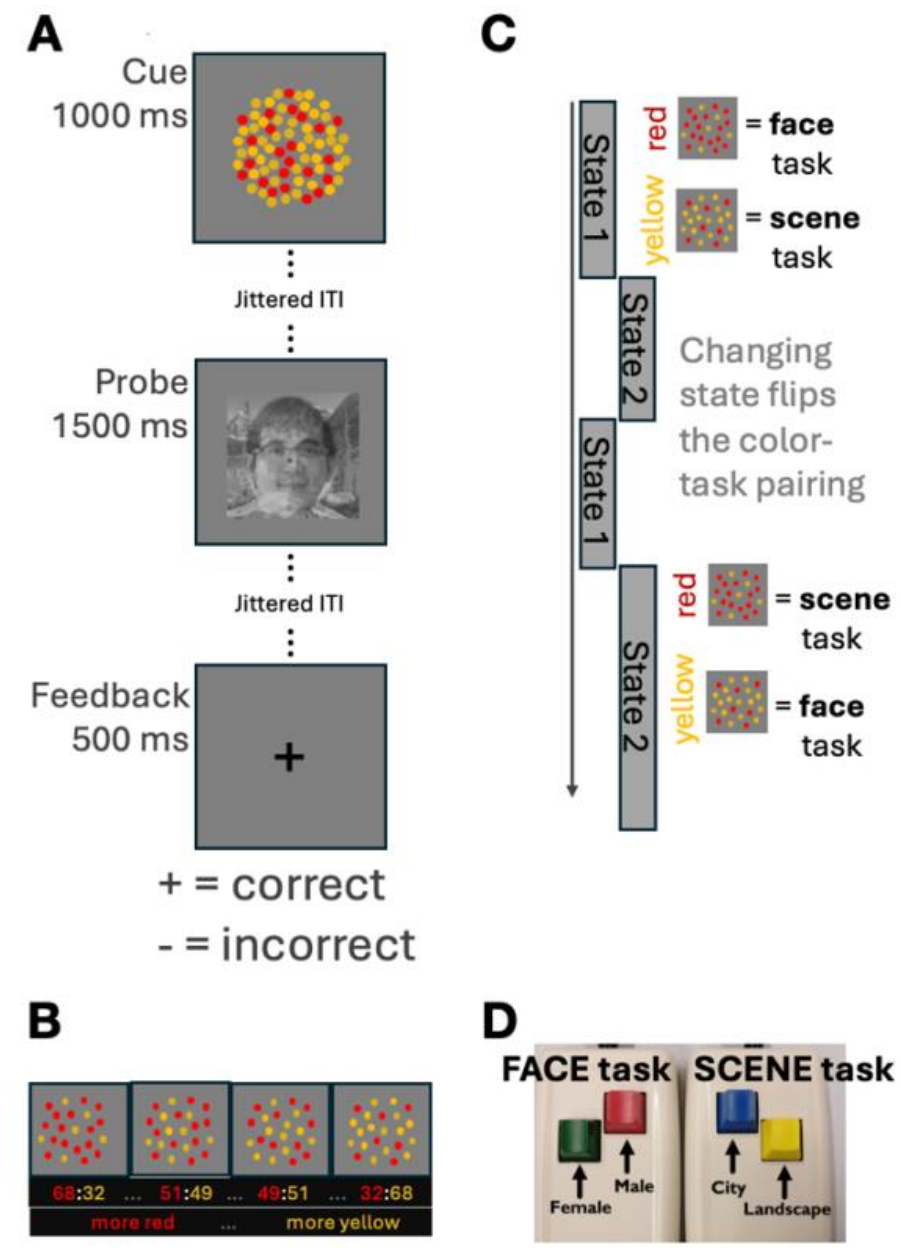
Experimental paradigm and Bayesian computational model. (A) Trial sequence. Participants integrated sensory evidence with an internally maintained contextual state to determine whether to perform a face or scene judgment. (B) Sensory evidence was manipulated by varying the proportion of red and yellow dots, producing graded perceptual uncertainty. (C) Hidden-state transitions reversed the mapping between cue color and task every 10–20 trials, requiring continual updating of contextual beliefs. (D) Subjects use different hands to indicate different task responses. The face image in this (A) is the senior author of this paper, included with permission.

Behavior was modeled using a Bayesian computational model that formalized this integration process (Leach et al., 2025, also see supplementary methods). On each trial, the model estimated a probabilistic color belief from the sensory evidence and a probabilistic state belief representing the current hidden cue-task mapping. These two beliefs were combined into a joint probability distribution, from which the model derived a trial-wise task belief corresponding to the inferred probability that the face or scene task should be performed. Following feedback, state beliefs were updated according to Bayesian inference, producing trial-by-trial changes in task belief. For the present study, we used trial-wise estimates of task belief as the primary moderator variable for BSM analyses. We additionally derived an uncertainty-weighted error signal by combining task belief with feedback outcome to quantify the extent to which feedback confirmed or contradicted participants’ inferred task representation. For each participant, source-region beta series and trial-wise computational moderators were standardized to zero mean and unit variance before constructing the interaction regressor.

We used task belief as the moderator variable for empirical validation because it generates a strong *a priori* prediction regarding motor-system recruitment. From the computational model, positive task-belief values correspond to the face task, whereas negative values correspond to the scene task. These task identities were associated with distinct motor effectors, specifically with face tasks performed using the left hand and scene tasks performed using the right hand (Fig 2D). Consequently, increasing positive task-belief values should be associated with stronger coupling with the right motor cortex, whereas increasingly negative task-belief values should be associated with stronger coupling with the left motor cortex. Because this prediction follows directly from the task structure and computational model, it provides a validation test for the BSM connectivity method.

In addition to task belief, we performed a second analysis using an uncertainty-weighted error signal derived from the computational model. We developed this variable in our previous model-based fMRI study (Leach et al., 2026). This variable combines trial-wise task belief with the observed feedback outcome to quantify the extent to which feedback confirms or contradicts the participant’s inferred task representation. Feedback that contradicts a strongly held task belief produces larger uncertainty-weighted errors than feedback received under uncertain task beliefs, reflecting greater evidence that the current internal representation should be updated.

Importantly, both task belief and uncertainty-weighted error are continuous variables derived from the computational model that change trial-to-trial, rather than discrete, block-alternating experimental conditions. They vary from trial to trial and correspond to distinct stages of the task, with task belief indexing the inferred task representation during the cue epoch and uncertainty-weighted error indexing feedback-related processing during the feedback epoch. This temporal separation provides an additional validation of BSM because they require connectivity to be estimated from continuously varying computational variables that are expressed at different task epochs.

For each participant, trial-wise BOLD response amplitudes were estimated using the same LSS procedure described above. Single-trial response amplitudes were estimated using LSS regression applied to BOLD time series after nuisance regression, including motion-related effects, without removing task-evoked responses. Specifically, we included the following nuisance regressors: rigid-body motion estimates (three translations and three rotations), cerebral spinal fluid, and white matter noise components from the component-based noise correction procedure (Behzadi et al., 2007). We also included derivatives and squares of these components. Error trials were retained and were not modeled separately during single-trial beta estimation.

A dorsal medial prefrontal cortex (dmPFC) region was selected as the seed because our previous analyses identified it as a frontoparietal region involved in representing integrated color and contextual information. The dmPFC seed region was defined based on our previously reported analysis identifying regions that responded to the integrated uncertainty of state and color information (Leach et al., 2025), thus independent of the task-belief and error signals we used as moderators in the present study. For each trial, task-belief or uncertainty-weighted error values were entered as the moderator variable, and BSM interaction coefficients were estimated between the seed and every voxel, akin to a seed-based functional connectivity approach, yielding whole-brain maps of trial-wise connectivity modulation. Separate analyses were performed for cue-, probe-, and feedback-related beta estimates to evaluate the temporal specificity of the connectivity effects. Individual BSM maps were entered into a second-level group analysis to identify brain regions exhibiting consistent task-dependent connectivity modulation across participants, by performing group-level t-test against zero. Whole-brain statistical maps were thresholded at a voxel-wise threshold of p < 0.005, followed by cluster-level correction for multiple comparisons (p < 0.05) using Monte Carlo simulations implemented in AFNI’s 3dClustSim. Spatial autocorrelation parameters were estimated from task-model residuals and averaged across participants. Cluster simulations used a non-Gaussian spatial autocorrelation function (ACF) model, with parameters of 0.67, 1.39, and 3.50. The resulting minimum cluster extent was 128 voxels (2 × 2 × 2 mm voxel size; 1,024 mm³). To assess whether the observed effects reflected changes in functional connectivity rather than regional response amplitude alone, we performed complementary whole-brain parametric modulation analyses using the same trial-wise moderator variables. This analysis tested whether the uncertainty-weighted error explained variability in evoked BOLD responses beyond the average event-related response and provided a specificity control for the BSM findings.

### 2.6 Data and code availability

Code and data are available at https://github.com/HwangLabNeuroCogDynamics/BSM and https://openneuro.org/datasets/ds006821.

## 3. Results

### 3.1 Statistical power of BSM

A useful task connectivity method should reliably detect task-dependent changes in functional connectivity when such effects are present. Therefore, we first evaluated the statistical power of BSM to detect task-modulated connectivity across a range of simulated interaction strengths and SNR (Fig 3). Power increased systematically with the magnitude of the simulated interaction effect, approaching ceiling levels for moderate-to-large interaction strengths (δ ≥ 0.20) across most SNR conditions. In contrast, when no interaction effect was present (δ = 0), significant interaction estimates were rarely observed. We explicitly tested for false positives in a later section. Power was influenced by both latent neural and BOLD signal quality (SNR). Across interaction strengths, higher latent SNR and higher BOLD SNR both improved detection rates.

**Figure 3.**
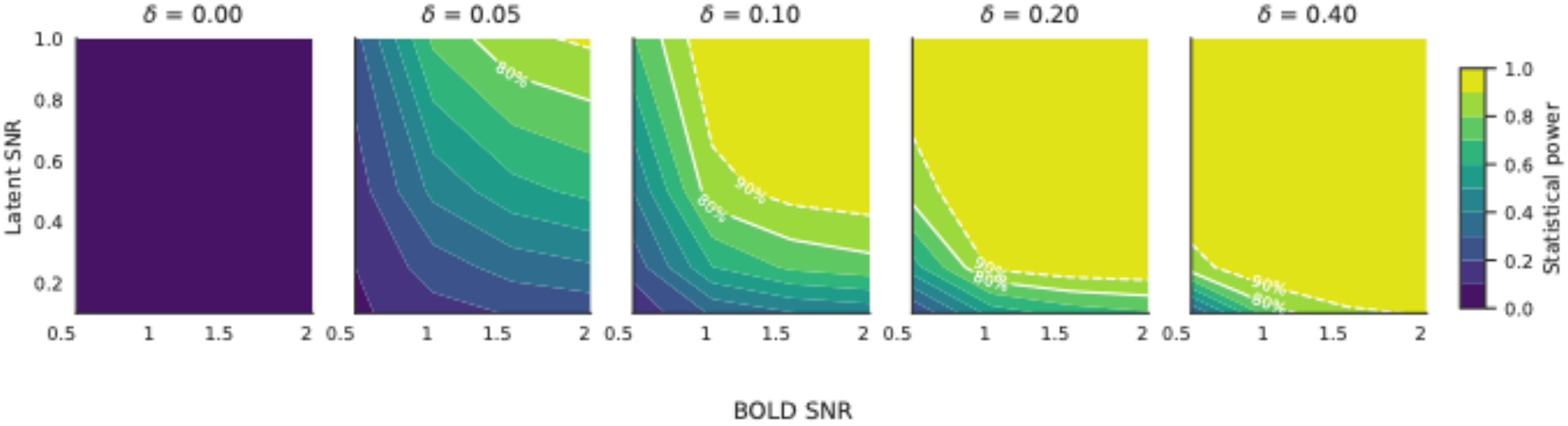
Statistical power of BSM across interaction strengths and SNR conditions. Color maps show the proportion of simulated experiments in which the interaction effect was detected at the group level (*p* < 0.05). Columns correspond to the ground-truth interaction strength (δ = 0, 0.05, 0.10, 0.20, and 0.40). Within each panel, the x axis denotes BOLD SNR and the y axis denotes latent neural SNR. White solid and dashed contour lines indicate 80% and 90% statistical power, respectively.

For small interaction effects (δ = 0.05–0.10), power increased substantially as BOLD SNR increased, with the highest detection rates observed when both latent and BOLD SNR were high. On the other hand, for small interaction effects, when latent SNR are low detection rate suffered even with relatively large BOLD SNR. At larger interaction strengths (δ = 0.20–0.40) power approached ceiling across SNR conditions. While ground truth latent SNR cannot be empirically measured with fMRI, using existing empirical data we found that most subcortical and cortical regions have SNR ranging from 1 to 2.5 (Chen et al., 2024; Leach et al., 2025). These results indicate that BSM can reliably detect task-dependent connectivity modulation under realistic signal conditions, with sensitivity increasing predictably as interaction effects become stronger and SNR increases.

### 3.2 Sensitivity to interaction strength

We next evaluated whether BSM interaction estimates systematically tracked changes in the magnitude of simulated connectivity modulation. Figure 4 shows the relationship between the ground-truth interaction parameter (δ) and the estimated standardized interaction coefficient across latent neural and BOLD SNR conditions. Across all conditions, estimated interaction coefficients increased monotonically with δ, demonstrating a consistent dose–response relationship between the simulated and estimated interaction effects. This relationship was approximately linear, with steeper slopes observed at higher latent neural and BOLD SNR. These findings indicate that BSM estimates are sensitive to changes in the underlying connectivity modulation, with greater sensitivity under higher signal quality conditions.

**Figure 4.**
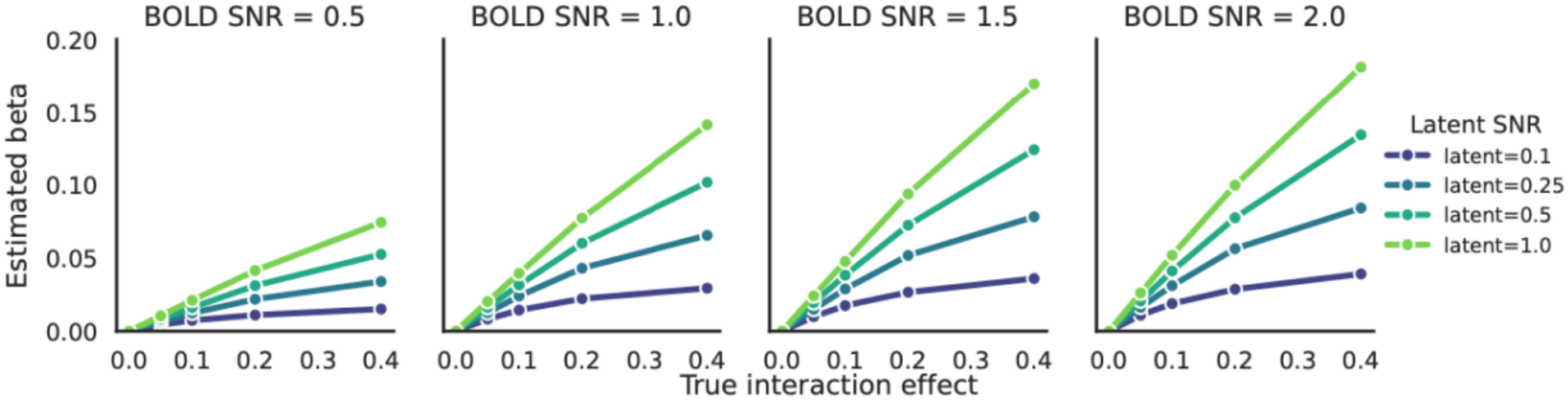
Recovery of simulated interaction effects by BSM. Mean estimated interaction coefficients are plotted as a function of the ground-truth interaction effect. Panels correspond to different levels of BOLD SNR, and colored lines indicate latent neural SNR.

### 3.3 Robustness to Baseline Coupling and Task-Evoked Coactivation

A primary motivation for the BSM method is to distinguish modulation of connectivity from other sources of variance that commonly occur in task-based fMRI experiments. In particular, strong intrinsic, task-independent coupling between regions and task-related coactivation can both produce signal correlations in regional activity, potentially complicating interpretation of connectivity estimates (Cole et al., 2019). To evaluate the specificity of the BSM interaction coefficient, we performed simulations that systematically varied baseline coupling, source evoked responses, correlations between source activity and the moderator, independent of the ground-truth interaction parameter.

We first systematically varied the magnitude of task-evoked source activation and baseline coupling while holding signal quality constant (latent SNR = 0.5, BOLD SNR = 1.0). Under this simulation configuration, noise was scaled proportionally to signal amplitude, such that changes in source activation magnitude did not substantially alter the correlation between source activity and the moderator. We found that interaction estimates were relatively stable across both source-effect and baseline-coupling conditions (Fig. 5A). For all combinations of task-evoked activation and intrinsic coupling, estimated interaction coefficients increased monotonically with the true interaction parameter, with relatively small differences across activation and baseline-coupling conditions. These findings indicate that BSM interaction estimates are relatively insensitive to changes in task-evoked activation magnitude and baseline coupling under the simulated fixed-SNR conditions.

**Figure 5.**
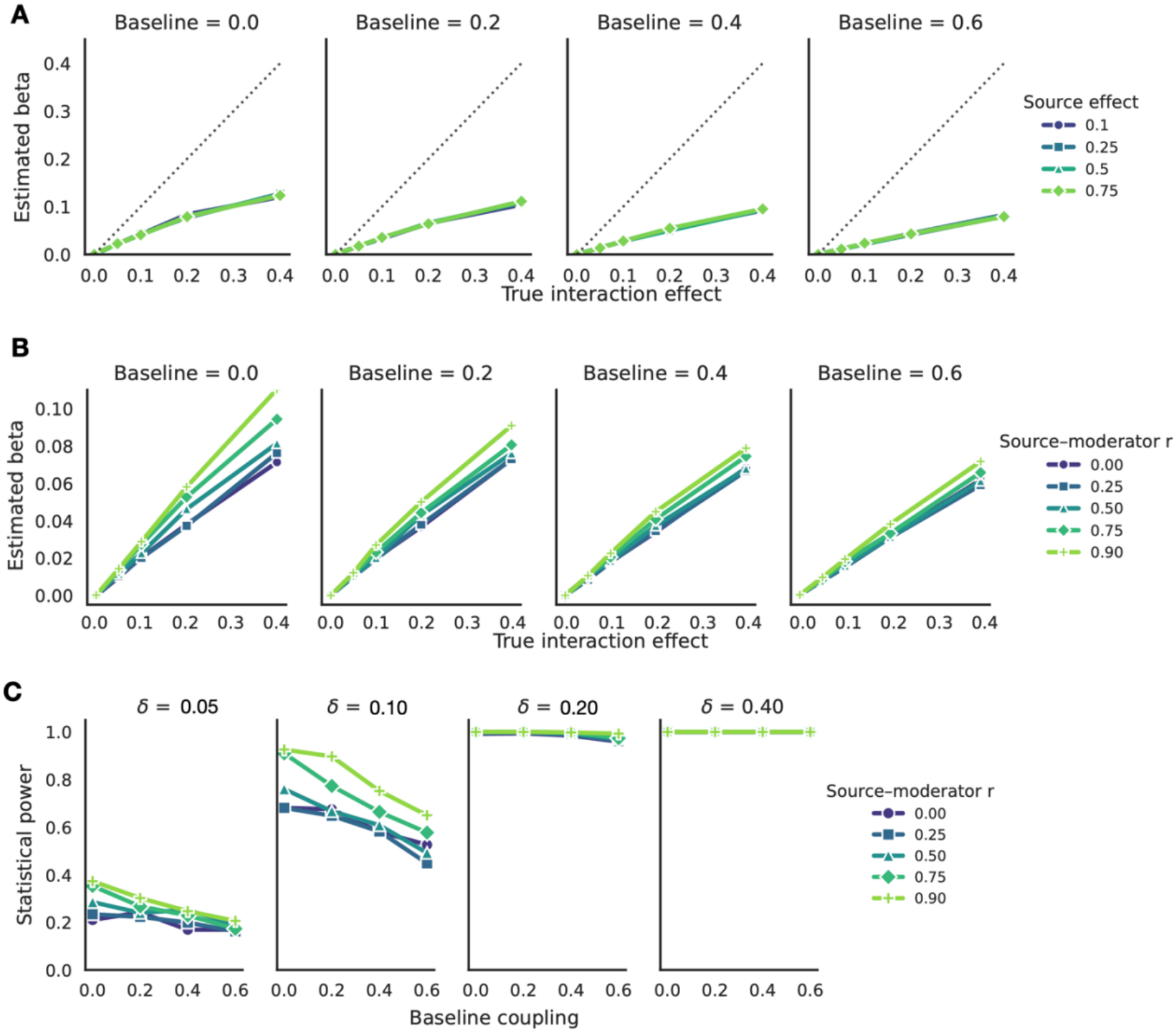
Influence of baseline coupling, task-evoked activation, and source–moderator correlation on standardized BSM interaction estimates. (A) Dose–response relationships between ground-truth interaction strength and estimated standardized interaction coefficients (beta) across baseline coupling and task-evoked activation (source effect) conditions. Panels correspond to different levels of baseline coupling, while colored lines indicate the magnitude of task-evoked source activation. Simulations were performed at latent SNR = 0.5 and BOLD SNR = 1.0. (B) Influence of source–moderator correlation and baseline coupling on standardized interaction estimates. Mean estimated interaction coefficients are plotted as a function of baseline coupling, with colored lines indicating different levels of source–moderator correlation. The ground-truth interaction strength ranged from δ = 0 to 0.4. Intended correlations ranged from 0 to 0.90. (C) Statistical power of BSM as a function of source–moderator correlation across interaction strengths. Statistical power was defined as the proportion of simulated experiments in which the interaction effect was significant at the group level (p < 0.05). For this simulation, baseline coupling was fixed at 0.2, with latent SNR = 0.5 and BOLD SNR = 1.0.

Because the correlation between source activity and the moderator may also influence interaction estimation, we conducted additional simulations that explicitly manipulated the correlation between the moderator and source signal, independently of baseline coupling between the source and target signals. We varied the latent source–moderator correlation from 0 to 0.90, baseline coupling from 0 to 0.60, and ground-truth interaction strength from 0 to 0.40. Unlike the above simulation, increasing source–moderator correlation systematically increased the magnitude of standardized interaction coefficients, whereas increasing baseline coupling generally reduced their magnitude (Fig. 5B). Statistical power also varied with source–moderator correlation and baseline coupling, with greater sensitivity at higher correlations and lower baseline coupling for small-to-moderate interaction effects (Fig. 5C). Collinearity between the interaction regressor and main effects remained minimal (Variance Inflation Factor = 1.01).

### 3.4 False positive controls

Another critical test is whether BSM can obtain spurious connectivity estimates when no interaction is present. To evaluate this possibility, we repeated the simulations with the ground-truth interaction parameter set at zero and quantified the proportion of significant interaction effects detected by BSM. Baseline coupling and task-evoked source activation were systematically varied across a wide range of values while all other parameters for the simulation remained constant. We found that false-positive rates remained close to the nominal α = 0.05 threshold across all conditions (Fig 6). Neither increasing baseline coupling nor increasing task-evoked source activation produced systematic elevations in false-positive rates. We also evaluated false positive control in the additional simulations that independently manipulated source– moderator correlation and baseline coupling (see section 3.3 above). False positive rates also remained around the 5% threshold across the full range of source–moderator correlations (minimum: 4.1%, maximum 5.3%,), with no systematic inflation as either parameter increased.

**Figure 6.**
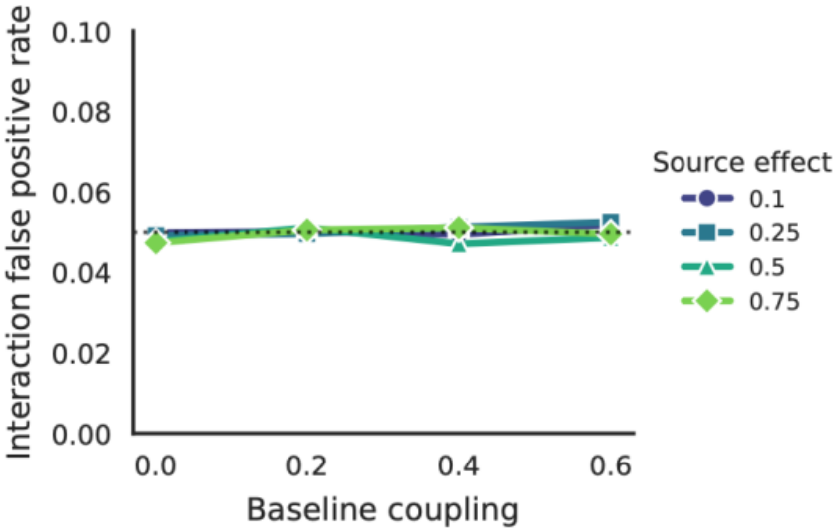
False-positive rates remain stable across baseline coupling and task-evoked responses strengths. Simulations were performed with the ground-truth interaction effect fixed at zero. The y-axis shows the proportion of simulations yielding a significant interaction effect, and the x-axis shows the magnitude of baseline coupling between source and target regions. Colored lines indicate the magnitude of task-evoked activation (moderator main effect). The dashed horizontal line denotes the nominal significance threshold (α = 0.05).

We further examined whether temporal dependencies in the moderator influenced false-positive control. In addition to the original independent Gaussian moderator, we simulated an autocorrelated Gaussian moderator using an AR(1) process (φ = 0.8) and a block-like moderator that persists for 10–20 trials. False-positive rates remained close to the nominal 5% threshold across all three conditions (4.3%, 4.1%, and 4.1%, respectively). Together, these results indicate that BSM maintains appropriate Type I error control across conditions.

### 3.5 Influence of experimental design parameters

We next examined how experimental design parameters influence the statistical power of BSM. Because BSM operates on trial-wise response estimates, power is expected to also be influenced by the amount of data available for estimating single-trial amplitudes and interaction effects. Power increased systematically with both the number of subjects and the number of trials (Fig 7 A). However, increasing the number of trials produced substantially larger gains in power than comparable increases in sample size. This effect was particularly strong for small-to-moderate interaction strengths, where increasing trial counts improved detection rates. We also evaluated the influence of temporal sampling rate (TR) and ITI jittering on performance (Fig 7B). Power improved as mean ITI intervals increased, reflecting improved estimation of trial-specific response amplitudes when events were less temporally overlapping. In contrast, variations in TR had comparatively modest effects on power.

**Figure 7.**
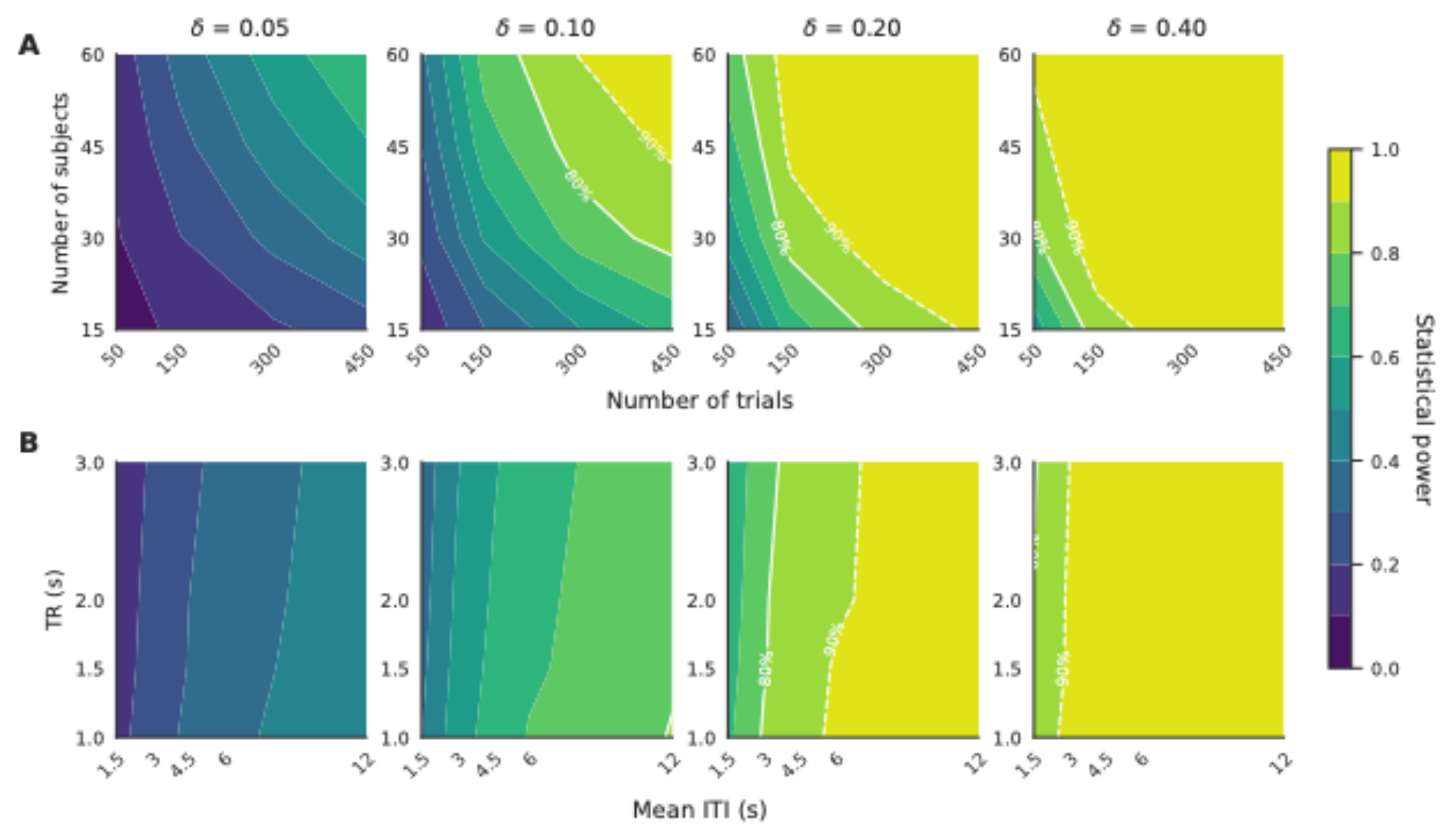
Influence of experimental design parameters on BSM statistical power. Color maps show the proportion of simulated experiments yielding a significant interaction effect. Columns correspond to different ground-truth interaction strengths (δ = 0.05, 0.10, 0.20, and 0.40). (A) Power as a function of the number of subjects (y axis) and the number of trials (x axis). (B) Power as a function of repetition time (TR; y axis) and mean inter-trial interval (ITI; x axis). Simulations were performed with latent SNR = 0.5 and BOLD SNR = 1.0.

### 3.6 Comparison with PPI

To benchmark BSM against an established task-connectivity approach, we compared its statistical power with PPI across the same simulation conditions. Both methods were evaluated using identical latent neural signals, hemodynamic model, and noise properties. For both methods, statistical power increased with interaction strength and SNR, approaching ceiling for strong interaction effects (Fig. 8A top row). Overall BSM and PPI exhibited comparable statistical power, averaging 78.1% and 77.1%, respectively, across interaction strengths SNR. However, their relative sensitivity varied with latent neural SNR. At lower latent SNRs (0.10 and 0.25), BSM exhibited modestly higher average power than PPI, by 4.1% and 2.3%, respectively. At higher latent SNRs (0.50 and 1.00), the methods showed nearly identical power or a modest PPI advantage, with differences of 0.2% and 2.2%. Both methods maintained false-positive rates around the 5% threshold (δ = 0; Fig. 8A bottom row).

**Figure 8.**
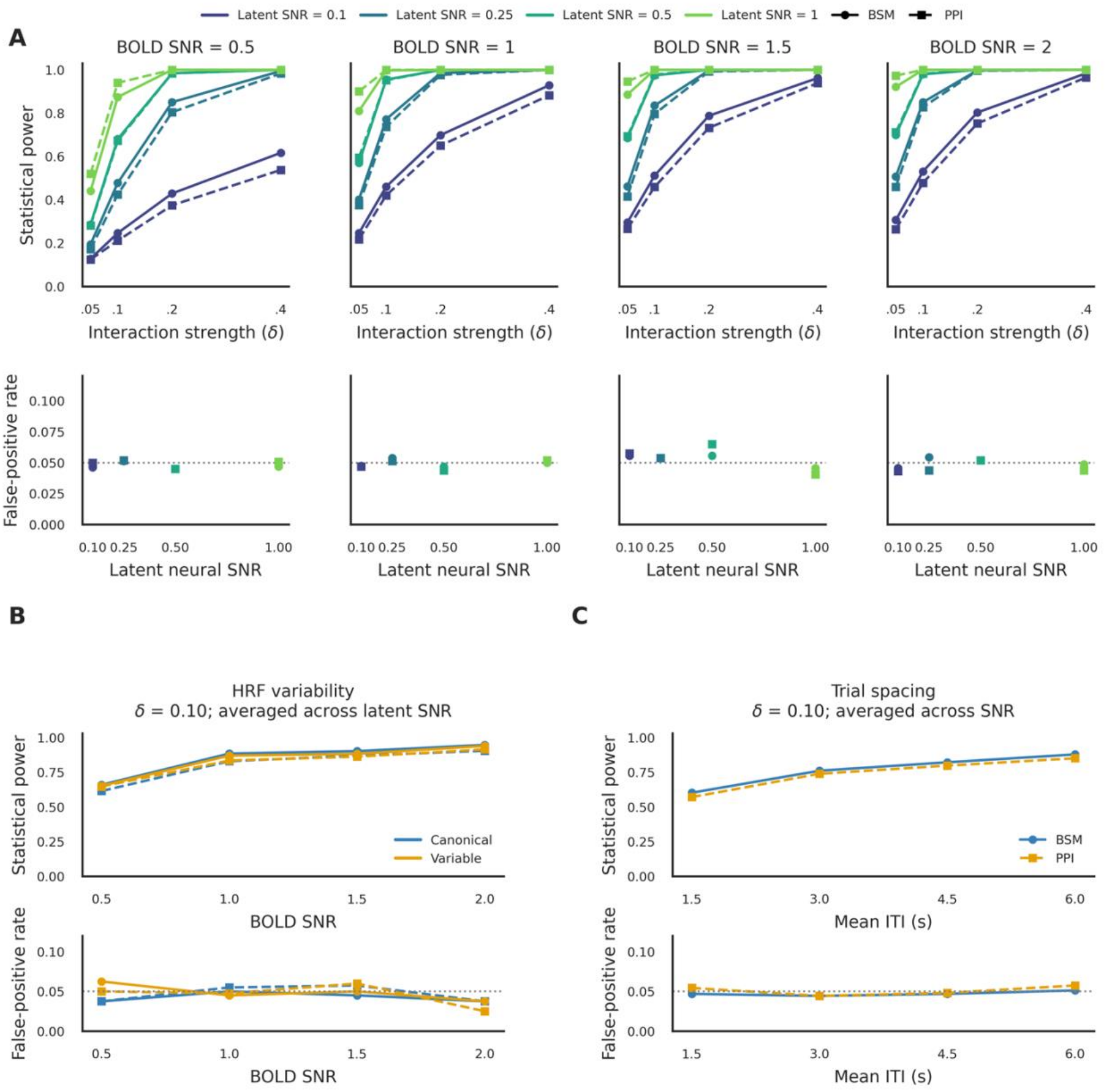
Comparison of statistical power and false positive rates between BSM and PPI. (A) Statistical power (top row) and false positive rates (bottom row) across BOLD and latent neural SNR conditions. Columns correspond to different BOLD SNR levels, and colors indicate latent neural SNR. In the top row, statistical power is plotted as a function of ground truth interaction strength (δ = 0.05–0.40). In the bottom row, false positive rates under the null condition (δ = 0) are plotted as a function of latent neural SNR. Results are averaged across task-evoked source activation and baseline coupling conditions, with mean ITI fixed at 4.5 s and variable HRFs. (B) Influence of HRF variability on statistical power (top) and false-positive rates (bottom) across BOLD SNR conditions. Colors distinguish canonical and variable HRFs. Statistical power is shown for δ = 0.10, averaged across latent neural SNR conditions, with source activation, baseline coupling, and mean ITI fixed at 0.5, 0.2, and 4.5 s, respectively. (C) Influence of mean ITI on statistical power (top) and false positive rates (bottom). Statistical power is shown for δ = 0.10, averaged across BOLD and latent neural SNR conditions, with source activation and baseline coupling fixed at 0.5 and 0.2, respectively. For all panels, solid lines and circular markers indicate BSM, whereas dashed lines and square markers indicate PPI. Horizontal dotted lines indicate the nominal false positive rate of 5%.

To examine whether statistical power depended on HRF variability, we conducted simulations contrasting the effects of canonical HRFs versus subject-specific HRF variability on statistical power (Fig. 8B). At an interaction strength of δ = 0.05, average statistical power was 84.9% for BSM and 80.5% for PPI under canonical HRFs, compared to 83.6% and 80.4% under variable HRFs. Thus, HRF variability had relatively small effects on statistical power for both methods. False-positive rates remained close to the 5% threshold under both HRF conditions. We further examined the influence of trial spacing by varying the mean ITI from 1.5 to 6 seconds (Fig. 8C). At δ = 0.10, increasing the mean ITI from 1.5 to 6 seconds increased statistical power from 60.3% to 87.9% for BSM and from 57.1% to 85.1% for PPI. This pattern indicates that both methods benefited similarly from longer trial spacing. The difference in statistical power between methods remained small across ITI conditions, ranging from approximately 2.2 to 3.1%. False-positive rates remained close to 5% across all ITIs for both methods. Together, these findings indicate that BSM and PPI exhibit broadly similar statistical sensitivity and appropriate false-positive control under our simulation.

### 3.7 Empirical Results

#### 3.7.1 Model-based functional connectivity analysis of task beliefs

We next evaluated BSM with empirical fMRI data. On each trial, participants inferred whether to perform a scene or face task by integrating perceptual inputs with an internally maintained state beliefs. These inferences were quantified using a computational model that generated a continuous task-belief estimates. Positive task beliefs indicated stronger belief that the current trial belonged to the face task, whereas negative values indicated stronger belief that it belonged to the scene task. The design provides a suitable test for the BSM analysis because task-belief estimates fluctuate probabilistically trial-by-trial as participants integrate evidence regarding the correct task rule (Fig 9A). In addition, the task structure generated a strong a priori prediction regarding motor-system activity. Face and scene tasks were associated with different response mappings, with face-task responses executed using the left hand and scene-task responses executed using the right hand. Because motor control is contralateral, increasing belief that the current trial belonged to the face task should be associated with stronger coupling between frontoparietal control regions and right motor cortex, whereas increasing belief that the current trial belonged to the scene task should be associated with stronger coupling with left motor cortex. Importantly, task-belief computations occur during cue processing, before participants make a response to the probe image or receive feedback during the feedback epoch.

**Figure 9.**
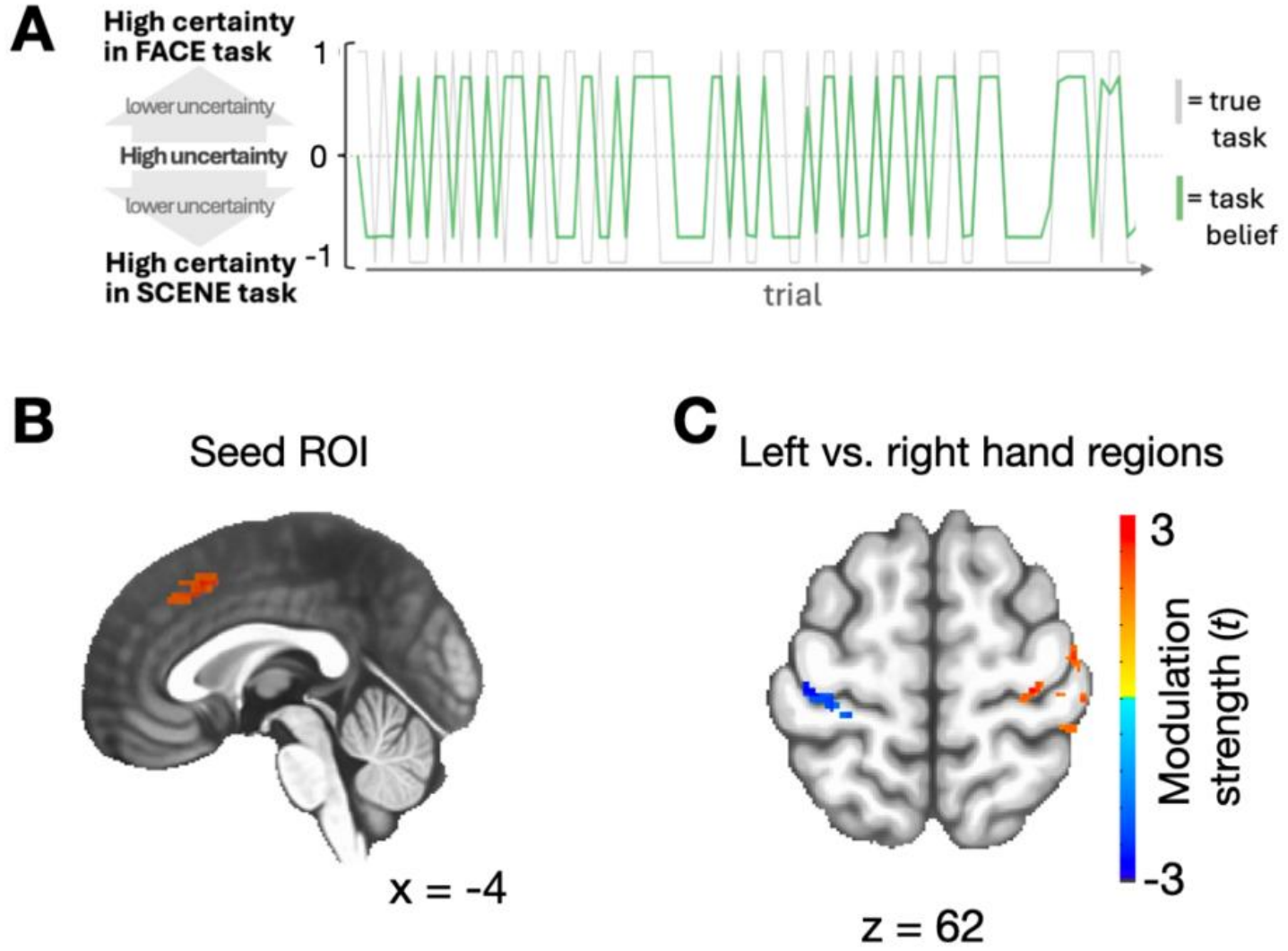
Empirical validation of BSM using trial-wise task-belief estimates. (A) Example trial-by-trial fluctuations in computationally derived task-belief estimates. Positive values indicate belief in the face task and negative values indicate belief in the scene task. (B) The frontal seed region used as the source ROI for whole-brain voxel-wise BSM analysis. (C) Connectivity modulation associated with task belief during the cue epoch. All brain maps thresholded at voxel level *p* < 0.005 and corrected cluster-level p < 0.05.

We therefore performed the BSM analysis restricting to cue-epoch’s single trial responses (Fig. 2), allowing us to test not only whether BSM detects connectivity modulation by a continuous computational variable, but also whether it localizes this modulation to the appropriate stage of the cognitive process. Demonstrating cue-specific connectivity would provide evidence that BSM can resolve trial-wise changes in functional interactions with the specificity afforded by event-related experimental designs.

We selected a medial dorsal prefrontal region as the source region of interest (Fig 9B). This region was selected based on our prior study (Leach et al., 2025), in which this clustered showed robust task-evoked response to the integrated uncertainty combining sensory evidence and internal state beliefs (cluster size = 955 2 mm^3^ voxel; MNI: x = -4, y = 22, z = 44). Using this ROI as the source region and performed BSM with every other voxel (akin to a seed-based functional connectivity approach), BSM indeed identified significant task-belief-dependent connectivity modulation with bilateral motor cortex during the cue period (Figure 9C). Positive task-belief values, corresponding to stronger belief in the face task, were associated with increased connectivity between the frontal seed and right motor cortex. Conversely, negative task-belief values, corresponding to stronger belief in the scene task, were associated with increased connectivity between the frontal seed and left motor cortex. These findings demonstrate that BSM can recover theoretically predicted patterns of task-modulated connectivity using a continuous moderator derived from a computational model.

#### 3.7.2 Model-based functional connectivity analysis of error signals

We next evaluated BSM using an uncertainty-weighted prediction error signal derived from the computational model. This variable incorporates the uncertainty of the inferred task belief, assigning larger values to feedback that contradicts a strongly held task representation. Such trial-wise variables are difficult to accommodate with conventional task-based connectivity approaches because they vary continuously from trial to trial. In addition, these modulations should be specific to the feedback period, not the cue or probe period. This trial epoch specific modulation test is relatively difficult to implement with other existing task functional connectivity methods.

As expected, conventional parametric modulation analysis revealed widespread regional BOLD responses associated with uncertainty-weighted errors, including distributed frontoparietal and posterior cortical regions (Fig. 10, left). In contrast, BSM identified a more selective pattern of connectivity modulation. Larger uncertainty-weighted errors were associated with stronger functional connectivity between the medial frontal seed region and distributed frontoparietal regions, as well as subcortical structures including the caudate and medial thalamus (Fig. 10, right). These connectivity effects were observed only when BSM was performed using feedback-epoch responses, whereas corresponding analyses of cue and probe epochs yielded no significant effects. Together, these findings demonstrate that BSM captures trial-wise modulation of functional interactions that is distinct from regional activation and localized to the appropriate cognitive stage of the task. Importantly, the connectivity findings cannot be explained by evoked responses within the source region alone. Conventional parametric modulation analysis identified regions whose evoked responses scaled with uncertainty-weighted errors, whereas BSM identified brain regions whose functional coupling with the medial frontal seed systematically varied with the same computational variable. Thus, BSM provides information beyond regional activation byrevealing how uncertainty-weighted errors modulate connectivity between distributed brain regions rather than simply modulating activity within individual regions.

**Figure 10.**
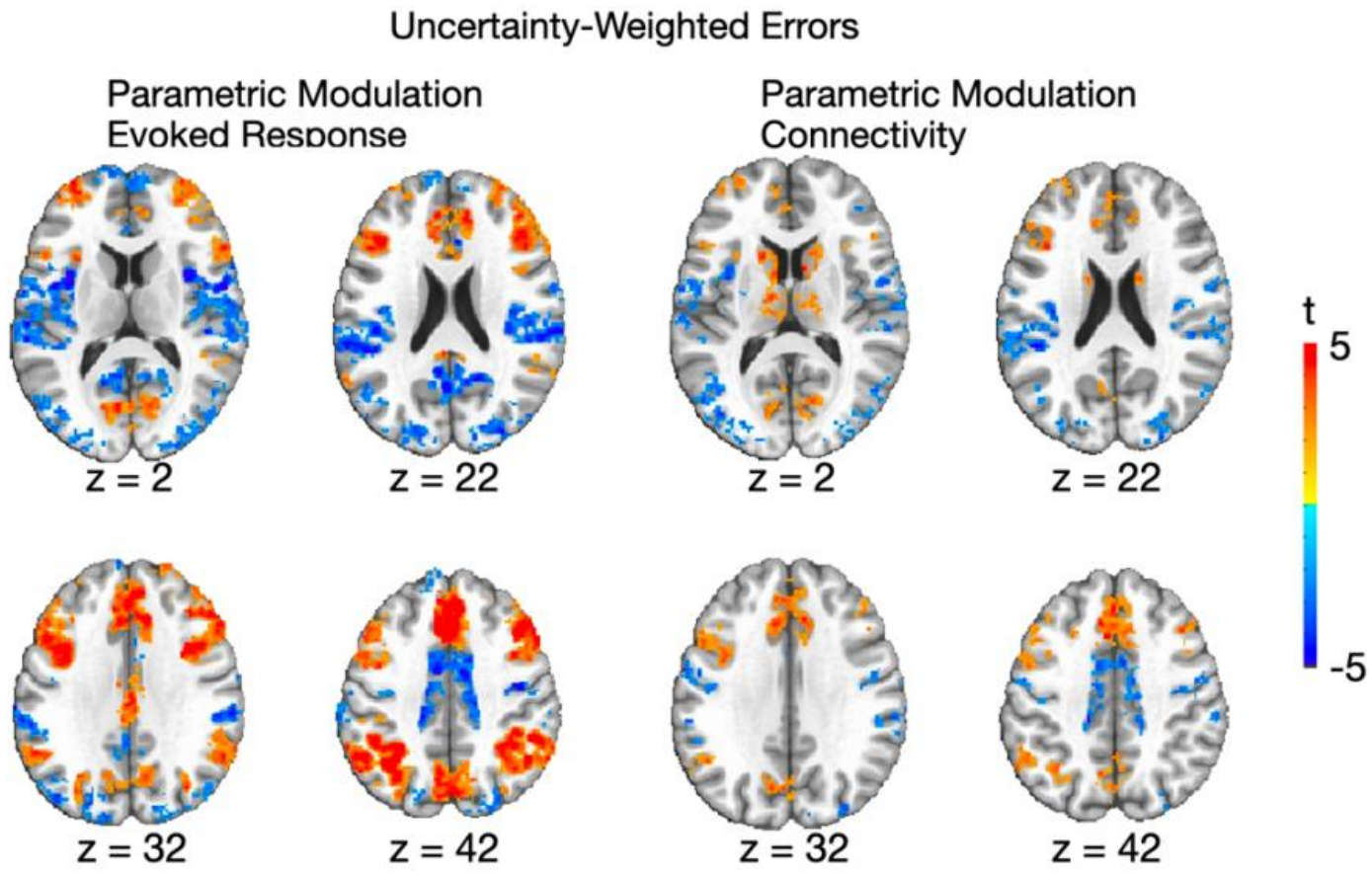
Regional activation and functional connectivity associated with uncertainty-weighted errors. Parametric modulation analysis of trial-wise uncertainty-weighted errors (left). BSM analysis using the same computational variable revealed selective modulation of functional connectivity (right). Statistical maps are thresholded at voxel-level *p* < 0.005 and cluster-level *p* < 0.05 (corrected).

To summarize, observations from the empirical validation demonstrated the effectiveness and trial specificity of BSM. The two computational variables exhibited dissociable patterns of connectivity modulation that were consistent with their hypothesized computational roles. Task belief modulated functional connectivity during the cue epoch, when the inferred task representation was used to prepare for subsequent action selection, but did not significantly modulate connectivity during the feedback epoch. Conversely, uncertainty-weighted errors selectively modulated connectivity during the feedback epoch, when outcome information was available, but not during the cue or probe epochs. In addition to trial-epoch specificity, the validation analyses were performed using the same medial frontal ROI as the source region. Thus, the observed effects further reflect selective changes in how the same frontal region communicate with distributed brain regions as cognitive demands evolve over the course of a trial. This flexibility illustrates a key advantage of BSM, allowing multiple model-derived variables to be tested across distinct trial epochs within an event-related fMRI experiment.

## 4. Discussion

Event-related fMRI combined with computational modeling has become a powerful approach for investigating the latent cognitive processes underlying human behavior. Although model-based fMRI is widely used to relate regional brain activity to trial-wise computational variables, comparatively few methods can test how these variables influence interactions between distributed brain regions. The present study addresses this methodological need by evaluating a regression-based BSM framework for estimating task-modulated functional connectivity using single-trial response amplitudes. By directly modeling the interaction between regional activity and task-related variables, this approach provides a flexible statistical framework for examining connectivity modulation by both categorical and continuous variables in event-related fMRI.

### 4.1 BSM is an effective task-based functional connectivity method for event-related fMRI

We showed that across a comprehensive set of simulations, BSM exhibited several desirable properties. The method can sensitively detect simulated interaction effects, maintained appropriate false-positive rates, and was relatively robust to changes in intrinsic coupling and task-evoked coactivation. Furthermore, these statistical properties translated to empirical data.

Using computational variables derived from a Bayesian model, BSM identified patterns of connectivity modulation associated with both task belief and error, feedback processes. Together, the simulation and empirical results demonstrate that regression-based BSM provides a practical approach for investigating how trial-wise variables modulate functional connectivity in event-related fMRI experiments.

The simulation results also provide insight into the conditions under which BSM is expected to perform well. Unlike PPI approaches that estimate interactions from continuous BOLD or latent neural time series, BSM operates on trial-wise response amplitudes. Because beta-series estimates serve as the inputs to the connectivity model, the precision of interaction estimates depends directly on the quality of the single-trial response estimates. Thus, improvements in latent neural and BOLD SNR both increased statistical power because they improved the estimation of trial-wise response amplitudes. Likewise, increasing the number of trials improved performance by providing more observations for estimating the interaction effect, whereas increasing ITIs reduced temporal overlap between neighboring hemodynamic responses and improved the precision of single-trial estimates. These observations have practical implications for study designs. Specifically, this suggests that event-related designs with large number of trials and adequate temporal separation between events are well suited for BSM.

Our simulations further showed that BSM achieved performance comparable to that of PPI across a broad range of simulation conditions. Both methods exhibited increasing statistical power when using higher signal quality data, with broadly similar statistical sensitivity and false positive control across the simulated conditions. Moreover, HRF variability had relatively small effects on both methods. Our findings suggest that the two methods are both good approaches for examining task-dependent functional connectivity. PPI is well established for examining task-dependent connectivity using continuous BOLD signals and both categorical and parametric task manipulations (O’Reilly et al., 2012). In contrast, regression-based BSM provides an alternative for event-related experiments, particularly when examining trial-wise variables or distinct epochs within trials (Ollinger et al., 2001).

The empirical validation analyses further illustrate the effectiveness and the potential specificity of BSM. The validation showed that BSM can dissociate functional interactions associated with distinct computational processes within the same experiment. The two model-derived variables produced different patterns of connectivity modulation that were consistent with their expected functional roles, with effects localized to distinct trial epochs. Task belief modulated connectivity only during the cue epoch, when participants inferred the appropriate task before making a response, whereas uncertainty-weighted error selectively modulated connectivity during the feedback epoch, when outcome information became available for updating internal task representations. Importantly, these dissociable connectivity patterns were identified using the same medial frontal seed region. Rather than simply characterizing whether a brain region participates in task-dependent connectivity, BSM revealed how the connectivity profile of the same source region changed as different computational processes unfolded across a trial. The temporal dissociation illustrates the utility of combining single-trial response estimation with moderation regression for studying distinct cognitive processes within event-related fMRI experiments.

One import component of the method is that all analyses were performed using single-trial response amplitudes estimated with the least-squares separate (LSS) procedure. LSS provides trial-wise estimates for event-related fMRI, and has often been used for decoding or representation similarity analyses (Mumford et al., 2014; Turner et al., 2012). However, overlapping hemodynamic responses can affect the precision of single-trial beta estimates (Mumford et al., 2012). Furthermore, the precision of single-trial estimates depends on experimental design characteristics, particularly the temporal spacing and overlap of hemodynamic responses across trials (Abdulrahman & Henson, 2016; Visser et al., 2016).

Consistent with this prior work, our additional simulations showed that shorter ITIs reduced the correlation between ground-truth trial amplitudes and LSS estimates. This creates opportunities to improve connectivity estimation through alternative methods of estimating beta-series. Because regression-based BSM operates on single-trial response estimates, it is not restricted to LSS and can be implemented using other beta-estimation methods, including least-squares-all (LSA).

Other methods such as GLMsingle (Prince et al., 2022), for example, jointly estimate voxel-wise hemodynamic responses, optimize noise regressors, and apply regularization across trials to improve the reliability of single-trial beta estimates. Because our simulations showed that the statistical performance of BSM depends strongly on the fidelity of single-trial response estimation, future work that further improve beta-estimation methods should in turn improve parameter recovery and statistical power for BSM.

### 4.2 Limitations

An important consideration in interpreting the empirical results is the spatial overlap between regional activation and connectivity modulation associated with uncertainty-weighted errors (Fig. 10). Although these patterns share a similar spatial distribution, this overlap does not necessarily indicate that activation confounds connectivity estimates. Regional activation and connectivity modulation are not mutually exclusive, as the same brain regions may exhibit both changes in response amplitude and changes in functional coupling associated with a cognitive variable. Importantly, the BSM regression includes both source activity and the moderator as main effects, such that the interaction coefficient captures modulation of the source–target coupling beyond their main effect contributions. Our simulations further demonstrated that increasing source–moderator correlation did not systematically inflate false positive rates under the tested conditions. Nevertheless, these results do not exclude potential influences of model misspecification, such as nonlinear task-evoked responses.

An important limitation of our study concerns the level we implement the latent neural response simulations. Our simulations modeled latent neural events that generated BOLD responses without simulating the detailed biophysical processes, such as neuronal firing rate, local field potentials, synaptic coupling, that generate fMRI signals. Several previous validation studies of functional connectivity methods have employed more detailed neural-mass or biophysical models that explicitly represent synaptic dynamics, potentials, or circuit interactions (Cole et al., 2019; Masharipov et al., 2024). The goal of our simulation was to control the magnitude and interactions between events that give rise to measurable BOLD responses rather than modeling the precise neuronal mechanisms that generated those events. We are therefore agnostic to the exact circuit level physiology that can generate the significant interaction effects we simulated. Because the relationship between microscopic neural activity and the BOLD signal remains indirect (and controversial), they depend on assumptions regarding neurovascular coupling (Logothetis, 2008). We therefore chose a simpler latent generative framework that minimizes assumptions while retaining necessary control over important properties that BSM is designed to estimate. Nevertheless, a more definitive evaluation of BSM will likely come from experiments that simultaneously measure neural activity and HRF. Animal neuroimaging have made such studies feasible (David et al., 2008; Mandino et al., 2025; Xie et al., 2025). These methods could evaluate how accurately regression-based BSM recovers task-dependent neural interactions from fMRI measurements, ultimately provide a stronger biological validation than simulations.

Another related consideration in interpreting the comparison between BSM and PPI is the generative model used in our simulations. In our simulations, coupling between brain regions was defined at the level of trial-wise neural response amplitudes, and thus although both methods were evaluated using identical simulated BOLD signals, this formulation is more closely aligned with the BSM specification and may influence their relative sensitivity. An alternative approach would model interactions as continuous, time-varying modulation of neural activity within task epochs. However, doing so would require additional assumptions about within-trial neural dynamics and the temporal structure of coupling, which we do not know the ground truths and thus require additional assumptions. We considered this level of biophysical modeling beyond the scope of the present simulations. Accordingly, our findings should be interpreted as characterizing the relative statistical sensitivity of BSM and PPI under a trial-evoked response model, rather than establishing the superiority of either method across all possible neural generative processes.

It is important to emphasize that BSM measures statistical interactions between brain regions and therefore does not estimate the direction or causal influences. Similar to conventional PPI and beta-series correlation approaches, the estimated interaction coefficients quantify changesin functional coupling rather than causal connectivity. Questions regarding directed communication or causal mechanisms remain better addressed using approaches such as dynamic causal modeling or brain stimulation in combination with fMRI (Hwang et al., 2020; Iyer et al., 2022; Nee & D’Esposito, 2016).

### 4.3 Implementation suggestion and conclusion

Model-based fMRI, which uses computational models to analyze fMRI data, is shifting fMRI analyses from comparisons between experimental conditions toward trial-by-trial characterization of latent cognitive processes (O’Doherty et al., 2007). BSM is well aligned with this approach because it allows computational variables, as well as categorical variables, to be incorporated directly into a statistical model of functional connectivity. In practice, BSM can be implemented using a variety of connectivity analysis strategies. Similar to conventional seed-based functional connectivity analyses, investigators can define one or more seed regions based on hypotheses or independent ROIs, and test how trial-wise variables modulate connectivity between the seed and the rest of the brain or selected target regions. This is the approach we demonstrated with our empirical validation. Alternatively, the same framework can be extended to whole-brain or network-level analyses by estimating trial-wise connectivity between all pair-wise brain regions. This will result in a network-level description of how trial-wise variables modulate large-scale functional brain network interactions. In one of our previous studies (Leach et al., 2026), we adopted this network-level BSM approach to characterize how frontoparietal connector hubs leverage multiple distinct control signals to dynamically adjust its interactions with distributed brain regions. For both seed-based and network-based, appropriate cluster corrections can be applied to control for multiple comparisons (Cox et al., 2017; Zalesky et al., 2010).

The method is equally flexible with respect to the moderating variable. Although we focused here on trial-wise variables derived from computational models, the same framework is readily applicable to a broad range of trial-wise psychological or experimental variables. Moderators can be continuous or categorical, including trial-to-trial self-reports, experimental conditions, physiological measurements, or latent variables derived from computational models as demonstrated in our empirical validation. This flexibility allows regression-based BSM to accommodate conventional condition-based connectivity questions as well as more fine-grained analyses of trial-wise and within-trial cognitive processes (with the appropriate epoched-design).

More generally, because BSM operates directly on trial-wise response amplitudes, it naturally complements other single-trial analysis approaches. This offers an opportunity to integrate connectivity with other information. For example, single-trial response estimates are often used for of multivoxel pattern analysis (MVPA), representational similarity analysis (RSA), encoding models, and related methods for characterizing neural representations (Kriegeskorte et al., 2008). By operating on the same trial-level data, BSM provides a method in which regional activation, representational geometry, and functional connectivity can be examined using the same response estimates.

In conclusion, regression-based BSM provides a flexible statistical framework for testing task-modulated functional connectivity using single-trial response estimates. By integrating conventional moderation regression with beta-series connectivity analysis, this approach accommodates both categorical experimental manipulations and continuous computational variables. Given the widespread use of event-related fMRI for studying human cognition, BSM provides a practical approach for investigating how functional brain networks change with task demands and cognitive processes.

## Author contributions are as follows

Conceptualization: KH

Methodology: KH

Investigation: SCL, SES, KH, JJ

Visualization: SCL, SES, KH

Supervision: KH, JJ

Writing—original draft: KH

Writing—review & editing: KH, SCL, JJ

Authors declare that they have no competing interest.

Research reported here was supported by a National Institutes of Mental Health grants R01MH122613, R01MH140248, and the Iowa Neuroscience Institute. This work was conducted on an MRI instrument funded by S10OD025025.

AI-assisted tools were used to check spelling and grammar.

