## Supplementary Materials for "Estimating trial-wise modulation of functional connectivity using event-related fMRI"

In this supplementary material section, we describe the mathematical formulation of our computational model for the event-related fMRI dataset we used for empirical validation. Note that the model and results were originally reported in our two previously published papers (Leach et al., 2025, 2026). We edited and reproduced the supplementary method we reported in Leach et al., 2026 here. Note for consistency, the verbal and mathematical descriptions are identical to our previous papers.

### 1. Computational Model

We developed a Bayesian model to capture the computational processes underlying cognitive integration in our behavioral paradigm (Fig. S1). The model estimates two probabilistic representations, a hidden state belief and a perceptual (color) belief, which are then combined into a joint probability distribution. From this joint distribution, the model derives a probabilistic task belief specifying which task the participant should perform. In other words, the model generates trial-by-trial estimates of input representations, their integrated product (i.e., the joint distribution), the associated uncertainty (entropy), and the resulting task belief. These trial-wise estimates were then used in model-based functional connectivity analyses.

In the model (Fig. S1), for trial i, the probabilistic state belief ($S_i$, representing the two color–task mappings) and probabilistic color belief ($C_i$) are combined into a 2×2 joint distribution. This joint distribution is used to infer the probabilistic task belief ($T_i$), where Task 1 corresponds to the face task and Task 2 to the scene task (Eq. 1).

$$p(T_i \mid C_i, S_i) \qquad (1)$$

Here, state belief ($S_i$), color belief ($C_i$), and task belief ($T_i$) are latent variables. State and color beliefs are inferred from observable inputs: the color proportion ($D_i$) and the participant's response ($R_i$). Because state belief evolves over time, the previous state belief ($S_{i-1}$) influences the current state belief ($S_i$), which in turn influences the subsequent belief ($S_{i+1}$). To capture this temporal dependency, we implemented a trial-wise diffusion process (Eq. 2), applied at the start of each trial.

$$p(S_i) = p(S_{i-1}) * (1 - \theta_0) + 0.5 * \theta_0 \tag{2}$$

In other words, at the beginning of each trial, a confusion in the form of random guess (i.e., 0.5 probability) of the state is introduced to the previous state belief with a weight of $\theta_0$. By applying this theta weight to the state belief, the state belief is brought closer to 0.5. How much closer to 0.5 the state belief gets depends on the specific value of $\theta_0$. The range of possible change in state belief ranges from 0 (i.e., no confusion) to 0.5 (high confusion).

Color belief ($C_i$) was estimated from the observable color proportion ($D_i$) using a sigmoid function, see equation 3.

$$p(C_i | D_i, \theta_1, \theta_2) = \frac{1}{1 + e^{-\theta_1 * \log(\frac{D_i}{1-D_i}) + \theta_2}} \tag{3}$$

The two free parameters (θs) estimating this sigmoid function determined the (1) slope (accounting for color uncertainty/perception) and (2) intercept (accounting for color bias).

The model predicts the response by combining the term $p(T_i \mid C_i, S_i)$ with two additional free parameters, the participant's respective error rates for the face ($\theta_3$) and the scene tasks ($\theta_4$), to predict the responses ($R_i$). $R_i$ was coded as 0 for correct task and correct response or 1, which could mean either correct task but incorrect response or incorrect task and either correct or incorrect response. Taken the above steps together, for the true task $t_i$, the prediction of generating the correct response given the free parameters, the belief of state and color input [i.e., $p(R_i = 0|\theta s, S_i, D_i)$], is estimated in the following manner:

$$p(R_i = 0|\theta s, S_i, D_i) = (1 - \theta_{2+t_i})p(T_i = t_i|C_i, S_i)\, p(C_i|D_i, \theta_1, \theta_2)p(S_i) \tag{4}$$

Where 2+$t_i$ indicates the error rate corresponding to the true task (either face or scene task depending on the current true task). The prediction of committing an error [i.e., $p(R_i = 1|\theta s, S_i, D_i)$] can be obtained using $1 - p(R_i = 0|\theta s, S_i, D_i)$. This step of predicting $R_i$ despite $R_i$ being an observable variable was necessary to estimate the participant's respective error rates for the face ($\theta_3$) and the scene ($\theta_4$) tasks, which are necessary for trial-to-trial belief updating. By using the trial-wise values of the directly observable variables ($D_i$ and $R_i$), the generative model was inverted to infer latent variable distributions (Fig. S1A).

For the first trial, the prior is initialized as a uniform distribution. As the trials progress, the posterior of the previous trial becomes the prior for the current trial. Updating the prior was achieved by multiplying the likelihood ($p(R_i|\theta s, S_i, D_i)$) and prior (i.e., the posterior of the previous trial, $p(\theta s, S_i|D_1, \dots, D_{i-1}, R_1, \dots, R_{i-1})$) and dividing them by a normalization constant (Equation 5).

$$p(\theta s, S_i|D_1, \dots, D_i, R_1, \dots, R_i) \propto p(R_i|\theta s, S_i, D_i)\, p(\theta s, S_i|D_1, \dots, D_{i-1}, R_1, \dots, R_{i-1}) \tag{5}$$

As shown in Equation 5, the inference procedure estimates the probability distribution for each free parameter. In the implementation of the model, each probability distribution is coded as a vector representing the probability at different values. We tested the logistic function's slope parameter over a range of 1.5–5, based on model fits to independently collected pilot behavioral data (not reported here). The range of values tested for the logistic function intercept parameter was log(0.45/0.55) to log(0.55/0.45), corresponding to actual color biases up to a 55:45 (dominant:non-dominant color) split. Error-rate parameters for both the face and scene tasks were tested between 0 to 0.35, based on potential error rate ranges for each task. Finally, the range tested for the parameter associated with the diffusion of state information across trials was 0 to 0.9. This range was also selected based on model fits in a pilot sample not reported here. We then used the softmax function to select the optimal value for each of the five theta parameters for each participant.

The proposed model treats both state belief ($S_i$) and color belief ($C_i$) as probabilistic representations (Fig. S1B) for. As a result, the 2x2 joint probability distribution encoded the combined uncertainty of these two sources of inputs. Moreover, because the joint probability distribution encodes the combined uncertainty of the inputs, the output, Ti, that is generated from the joint probability distribution is also probabilistic (Fig. S1B). State belief must be maintained across trials and updated with feedback. Equations 4 and 5 specify this update, including theta parameters for the respective error rates for the face ($\theta_3$) and the scene tasks ($\theta_4$).

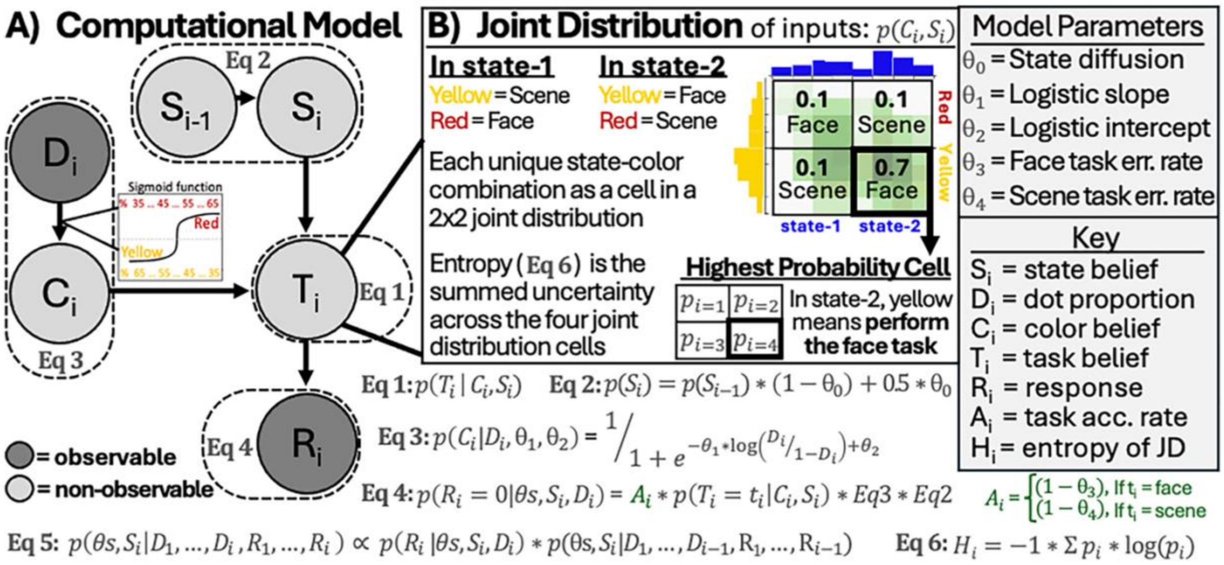

**Fig S1. Computational Model.** Note this is a reproduction of Fig 3 from Leach et al., 2025.
**(A)** Computational (Bayesian generative) model. Observable information ($R_i$ and $D_i$) was used to infer three latent variables: state belief (Si), color belief (Ci), and task belief (Ti). A sigmoid function (see Equation 3) was applied to the presented color proportion ($D_i$) to get an estimate of color belief ($C_i$). The state belief $S_i$ was inferred from $D_i$, and $R_i$. To allow the previous state to influence the current state, trial-wise diffusion of $S_i$ was implemented (see Equation 2). $S_i$ and $C_i$ were integrated into a 2x2 joint distribution from which the task belief $T_i$ was inferred (see Equation 1).
**(B)** In the example of a trial-level 2x2 joint distribution shown here, the probability that the face task was the task-to-perform is 0.8, summed from 0.1 (state = 1 and color = red) + 0.7 (state = 2 and color = yellow). Gray boxes on the far right side indicate the free model parameters fit during the inference procedure (see Equations 4-5) and a legend clarifying letter codes used in the model depiction and equations. These parameters include a diffusion parameter (allowing $S_{i-1}$ to influence $S_i$), a logistic function intercept and slope (capturing color bias and the high-uncertainty range, respectively), face and scene task error rates (used for updating following incorrect feedback; see Methods section 2.4 for full details).
